# Leptin regulates glucose homeostasis via the canonical WNT pathway

**DOI:** 10.1101/2021.02.16.431518

**Authors:** Kaj Kamstra, Mohammed Z. Rizwan, Julia A. Horsfield, Dominik Pretz, Peter R. Shepherd, David R. Grattan, Alexander Tups

**Author notes:** **Correspondence to:** Dr. Alexander Tups, Department of Physiology, University of Otago, PO Box 56, Dunedin, New Zealand. These authors contributed equally (Kaj Kamstra, Mohammed Z. Rizwan, David R. Grattan, Alexander Tups).

## Abstract

Leptin is a body weight regulatory hormone, but it is arguably even more potent at regulating blood glucose levels. To further our understanding of the molecular mechanisms by which leptin controls glucose homeostasis, we have used transgenic zebrafish models and conditional deletion of beta catenin in the mediobasal hypothalamus of adult mice to show that Wnt signalling in the brain mediates glucoregulatory effects of leptin. In zebrafish, under normal feeding conditions, leptin regulates glucose homeostasis but not adipostasis. In times of nutrient excess, we found that leptin also regulates body weight and size in this species. Using a Wnt signalling reporter fish, we show that leptin directly activates the canonical Wnt pathway *in vivo*. Pharmacological inhibition of this pathway prevented the leptin-induced improvement in glucose tolerance. In adult mice, conditional deletion of the key Wnt effector molecule, β-catenin, in the mediobasal hypothalamus of male mice confirmed the essential role of the Wnt pathway in mediating leptin action and the neuroendocrine regulation of glucose homeostasis. Adult-onset β-catenin deletion in the mediobasal hypothalamus led to glucose intolerance, exacerbation of caloric intake and body weight gain under high fat diet, as well as resistance to exogenous leptin.

## Introduction

The hormone leptin is known for its role in regulating energy balance. Although leptin is the primary adipostatic factor in mammals, it is well-established that leptin also regulates glucose homeostasis, independent of its adipostatic actions [1-5]: First, leptin is more potent at regulating glucose levels in blood than it is at suppressing appetite [6]. Second, acute disruption of leptin action *in vivo* raises blood glucose and plasma insulin levels before effects on body weight become apparent, and treatment of leptin-deficient Lep^ob/ob^ mice with leptin corrects glucose levels before body mass [7]. Third, Lep^ob/ob^ and leptin receptor-deficient Lepr^db/db^ mice become hyperinsulinemic before they become obese [8]. Fourth, humans who suffer from lipodystrophy, and rodent models of this disease, characterized by very low body fat and leptin levels, exhibit hyperglycemia, hyperinsulinemia and insulin resistance. All of these symptoms are corrected by leptin therapy [9, 10], which received approval by the FDA for this treatment purpose [11]. While leptin predominantly acts through the janus kinase 2 – signal transducer and activator of transcription 3 (JAK2-STAT3) pathway to regulate body weight [12-15], it seems that alternative pathways mediate the effect on glucose homeostasis [16, 17]. However, these pathways remain poorly defined.

Genome-wide association studies (GWAS) identified polymorphisms in several genes of the canonical Wnt pathway that increase the risk of glucose intolerance and type 2 diabetes (T2DM) [18-20]. The strongest effect size was associated with polymorphisms in the transcription factor 7-like 2 (*TCF7l2*) gene [21], which encodes a transcription factor of the canonical Wnt pathway. Wnt signalling is activated when a Wnt ligand binds to the frizzled (Fzd) receptor, which subsequently forms a complex with the co-receptor lipoprotein related protein (LRP) 5/6. This causes disheveled (Dvl) to phosphorylate LRP, which then inactivates glycogen synthase kinase 3β (GSK3β). GSK3β inactivation decreases phosphorylation of the transcriptional co-activator β-catenin. Stabilized β-catenin then enters the nucleus where it associates with transcription factors, such as TCF7l2, to ultimately regulate the transcription of downstream target genes [22]. Although canonical Wnt signalling has been studied extensively in the contexts of embryonic development and tumorigenesis, much less is known about its role in energy homeostasis [23]. Our laboratory showed that canonical Wnt signalling, specifically in the hypothalamus, is impaired during obesity and reinstated by leptin treatment [24]. Furthermore, we showed that GSK3β action specifically in the hypothalamus appears essential for glucose homeostasis. Lep^ob/ob^ mice were found to have elevated levels of active hypothalamic GSK3β, and glucose intolerance in these mice was acutely ameliorated by intracerebroventricular injection of a GSK3β inhibitor [25].

To test the hypothesis that leptin regulates glucose homeostasis via the canonical Wnt pathway, we decided to evaluate leptin action in a zebrafish model. Leptin signalling is evolutionarily well-conserved. Homologues for leptin and the leptin receptor are present even in invertebrate species like *Drosophila melanogaster* [26], and although leptin from species of different animal classes have low primary sequence homology, the secondary, tertiary, and quaternary structure, as well as key amino acids required for leptin’s physiological activity, are evolutionarily conserved [27]. Zebrafish (*Danio rerio*) express two leptin paralogues: leptin-a and leptin-b [28]. Both, like all vertebrate leptin paralogues, consist of four alpha helices, and contain a pair of cysteine residues that form a disulfide bridge. Three receptor interaction sites have been mapped, and each of these has at least some degree of amino acid sequence conservation [29]. Despite the conservation between species, it has been reported that leptin does not mediate adipostasis in zebrafish, but rather has an essential role in regulation of glucose homeostasis [30]. These data suggest that the glucoregulatory actions of leptin may, in fact, be the evolutionarily earlier function, with adipostasis added in higher vertebrates.

Here, we have used a transgenic Wnt-reporter zebrafish line to demonstrate that leptin activates canonical Wnt signaling, mediated via the leptin receptor, and that this pathway contributes to the glucoregulatory action of leptin. We subsequently tested whether this action is preserved in mammals, using conditional ablation of β-catenin in the mediobasal hypothalamus (MBH) of adult mice to prevent Wnt signalling. Mice lacking β-catenin in the MBH showed impaired glucose tolerance, and when on a high fat diet, they also showed markedly increased weight gain. These data demonstrate an essential role of the canonical Wnt pathway for mediating leptin action in the hypothalamus and show that this action contributes to the regulation of glucose homeostasis.

## Results

### Leptin activates the canonical Wnt pathway in vivo

To investigate whether leptin activates the canonical Wnt pathway, we used a transgenic zebrafish line (*Tg(7xTCF-Xla*.*Siam:nlsmCherry)*^*ia5*^) that sensitively detects translocation of the TCF7l2-β-catenin complex into the nucleus, thereby indicating canonical Wnt pathway activity [31]. Because the Wnt pathway is strongly active in embryonic patterning, we first established that at 5 days post fertilization (dpf), the developmental Wnt pathway activity has subsided to a level where it is mostly confined to the heart (figure 1A). The canonical Wnt pathway can be pharmacologically activated with lithium chloride (LiCl), which inhibits GSK3β [32], or inhibited with pyrvinium pamoate or PNU74654. Pyrvinium pamoate is an anti-helminthic drug that potentiates the activity of casein kinase 1α (CK1α), leading to enhanced degradation of β-catenin [33]. PNU74654 disrupts the interaction between β-catenin and TCF/LEF transcription factors [34]. We demonstrated that pharmacological activation or inhibition of the canonical Wnt pathway reliably increases or decreases the fluorescent signal in *Tg(7xTCF-Xla*.*Siam:nlsmCherry)*^*ia5*^ larvae in multiple tissues, particularly the heart but also in the hypothalamus (figure 1B,C). Strikingly, recombinant mouse leptin appeared to be efficacious in zebrafish, and treating 5 dpf *Tg(7xTCF-Xla*.*Siam:nlsmCherry)*^*ia5*^ larvae with leptin (100 nM) for 2 hours led to robust activation of the fluorescent construct specifically in the hypothalamus (figure 1D-F). In the hypothalamus, leptin increased fluorescence intensity significantly compared with vehicle-treated larvae, whereas in the heart leptin did not significantly increase fluorescence intensity.

**Figure 1.**
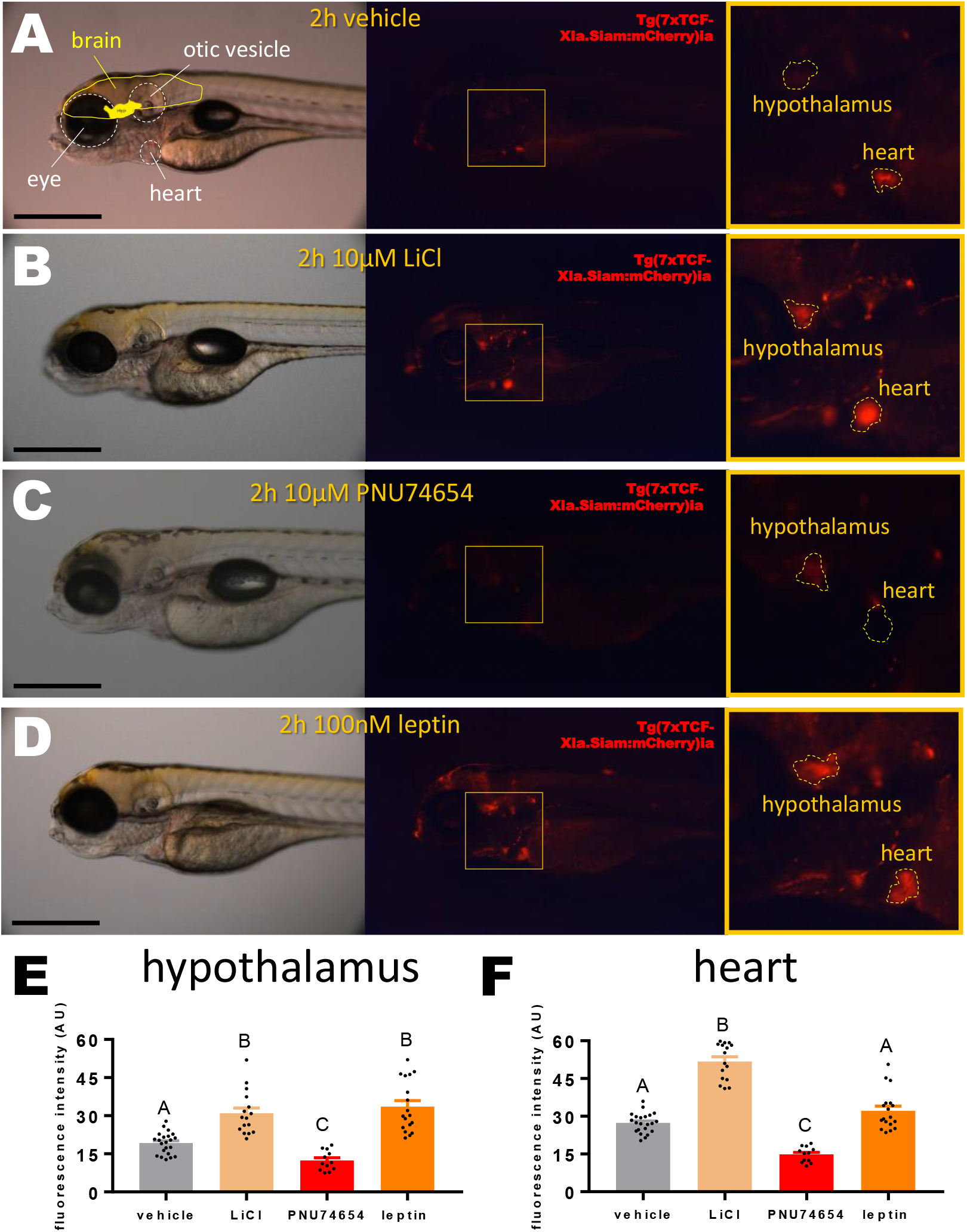
Wnt pathway activation by leptin in Tg(7xTCFXla.Siam:nlsmCherry)^ia5^larvae. (A) 5dpf Tg(7xTCFXla.Siam:nlsm Cherry)^ia5^ larvae treated with vehicle (Cortland salt solution). Left: bright field image, with anatomical landmarks encircled; Middle: Epifluorescence image; Right: Magnification of yellow box in middle image, with hypothalamus and heart encircled. Scale bar = 500 μM. (B) 5dpf Tg(7xTCFXla.Siam:nlsm Cherry)^ia5^ larvae treated with 10 μM LiCl for 2 hours. (C) 5dpf Tg(7xTCFXla.Siam:nlsm Cherry)^ia5^ larvae treated with 10 μM PNU74654 for 2 hours. (D) 5 dpf Tg(7xTCFXla.Siam:nlsm Cherry)^ia5^ larvae treated with 100 nM recombinant leptin for 2 hours. (E) Fluorescence intensity in the hypothalamus of differentially treated 5 dpf Tg(7xTCF-Xla.Siam:nlsmCherry)^ia5^ larvae. A-B=P<0.05, one-way ANOVA. (F) Fluorescence intensity in the heart of differentially treated 5 dpf Tg(7xTCF-Xla.Siam:nlsmCherry)^ia5^ larvae. Means ± SEM, A-B=P<0.05, one-way ANOVA.

### Leptin treatment ameliorates hyperglycemia in leptin deficient and wild type zebrafish

To further investigate the mechanism of leptin action on Wnt signaling, we created CRISPR-mediated knockout zebrafish lines on an AB_z_ background (figure S1), and conducted a series of studies to evaluate body weight and glucose homeostasis in lacking *leptin-a (lepa*^*nz301*^), *leptin-b* (*lepb*^*nz302*^) or the leptin receptor (*lepr*^*nz303*^). Raising the fish at identical tank densities, we found that body weight and standard length did not differ between wild type zebrafish and any of the knockout lines that were created (figure S2, S3), neither in males nor females at four, six or twelve months of age. To investigate whether leptin ameliorates hyperglycemia in leptin- or leptin receptor-deficient zebrafish, we induced a hyperglycemic state by immersing male and female zebrafish (n=5) in a 1% glucose solution for four days [35]. Immersion in 1% glucose steadily elevated basal blood glucose levels at a rate of 15-20 mg/dl per day, whereas immersion in normal system water did not change basal blood glucose levels (figure 2A-D). On the third day of immersion, one hour before blood sampling, fish were treated with either recombinant mouse leptin (2 mg/kg) or vehicle (Cortland salt solution). Leptin ameliorated hyperglycemia in wild-type and both leptin-a and leptin-b deficient zebrafish (figure S4), but not in leptin receptor-deficient zebrafish (figure 2C, D). Interestingly, the pattern of blood glucose elevation, and the effect of leptin on hyperglycemia was identical between males and females. Female zebrafish have a more variable body weight compared with males, due to the fact that they continuously produce eggs, which can make up to 25% of their total mass. For these reasons we performed all subsequent experiments in males only.

**Figure 2.**
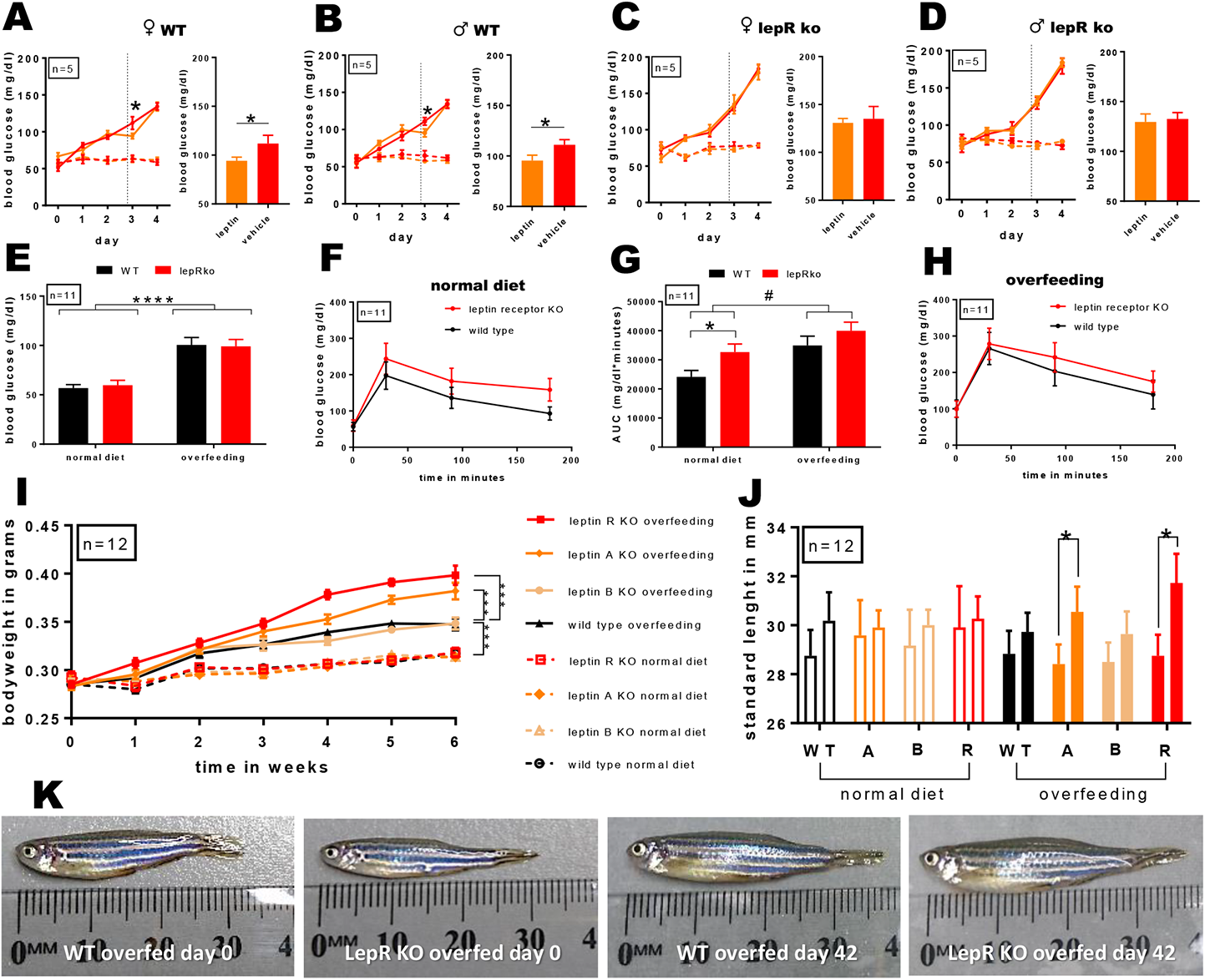
Regulation of glucose homeostasis and body weight in zebrafish. (A-D) Blood glucose values of male and female wild type and leptin receptor deficient “lepr^nz303^” zebrafish over time following immersion in a 1% glucose solution (left; solid lines = 1% glucose immersion, dotted lines = normal water immersion; *P<0.05, repeated measures ANOVA). On the third day, one hour before blood sampling, fish were injected intraperitoneally with recombinant mouse leptin (2mg/kg) or vehicle (right). Data displayed as mean±SEM. (E) Baseline blood glucose levels of lepr^nz303^ fish and wild type controls. ****P<0.0001, two-way ANOVA. (F) Glucose tolerance in lepr^nz303^ fish and wild type controls (n=12). (G) Area under the curve of (F) and (H). *P<0.05, one-way ANOVA; #P<0.05, two-way ANOVA. (H) Glucose tolerance in overfed lepr^nz303^ fish and wild type controls (n=12). (I) Body weights of lepa^nz301^ fish, lepb^nz302^ fish, lepr^nz303^ fish and wild type controls on a 6-week normal diet or overfeeding regime. ***P<0.001, repeated measures ANOVA. (J) Standard length of normal-fed and overfed lepa^nz301^ fish, lepb^nz302^ fish, lepr^nz303^ fish and wild type controls at the start (left) and end (right) of the feeding paradigm. *P<0.05, one-way ANOVA. (K) Examples of leptin receptor deficient and wild type zebrafish at the start and end of the overfeeding regime.

### Overfeeding reveals an effect of leptin on body size regulation in zebrafish

Our data are consistent with earlier studies suggesting that under normal feeding conditions, leptin regulates glucose homeostasis but not adipostasis in the zebrafish [30]. We next investigated whether leptin- and leptin receptor-deficient zebrafish were more prone to impaired glucose tolerance or diet-induced obesity (DIO). To this end, we exposed *lepa*^*nz301*^ fish, *lepb*^*nz302*^ fish, *lepr*^*nz303*^ fish, or wild-type control fish (n=12) to an overfeeding regime or a normal diet for six weeks. Glucose tolerance was tested at the start and end of this period. Intraperitoneal glucose tolerance tests (ipGTTs) revealed that although basal blood glucose levels were not significantly different (figure 2E), glucose clearance in leptin receptor knockout fish was reduced by 26% compared with wild type fish (figure 2F, G). Overfeeding increased basal blood glucose levels (from 64.1±3.5 to 100.5±1.8 mg/dl, P<0.001) and impaired glucose tolerance by 25%, independent of genotype (figure 2E, G, H). Surprisingly, we found that overfeeding also revealed an effect of leptin on body weight, with *lepr*^*nz303*^ (0.40±0.01 g, P<0.001) and *lepa*^*nz301*^ (0.38±0.01 g, P<0.001), but not *lepb*^*nz302*^ fish (0.35±0.01 g) having significantly increased body weight compared to overfed wild type controls (0.34±0.01 g; figure 2I). There was also an increase in standard length from 28.4±0.2 mm to 30.5±0.3 mm (P<0.05) in *lepa*^*nz301*^ fish and 28.8±0.25 mm to 31.7±0.36 mm (P<0.05) in *lepr*^*nz303*^ fish (figure 2 J, K). These results confirm that in zebrafish, under normal feeding conditions, leptin regulates glucose homeostasis but not body weight, consistent with the concept that this might be the evolutionarily conserved role of leptin. However, in times of nutrient excess, leptin did impact on both body weight and standard length.

### Activation of the canonical Wnt pathway and glucose lowering effects of leptin are dependent on a functional leptin receptor

Using the Crispr/Cas9 transgenic lines, we could investigate whether the glucoregulatory actions of canonical Wnt signalling are dependent on a functional leptin system. To confirm that Wnt reporter activation was mediated through leptin signalling, we used CRISPR/Cas9 to create *lepR* ‘crispants’ that are mosaic for leptin receptor knockout, and incubated them with either recombinant leptin or 10 μM LiCl (figure 3A) at 5 dpf. Mosaic knockout of the leptin receptor blocked the ability of leptin, but not that of LiCl to activate WNT signaling in the hypothalamus (figure 3B). We revisited the experimental paradigms described above to evaluate whether LiCl-induced activation of Wnt signalling could mimic the action of leptin in improving glucose tolerance, using *lepa*^*nz301*^ fish, *lepb*^*nz302*^ fish and *lepr*^*nz303*^ fish. Leptin treatment improved glucose tolerance in *lepa*^*nz301*^ fish (by 30%, figure 3C, F) and *lepb*^*nz302*^ fish (by 20% figure 3D, G). In *lepr*^*nz303*^ zebrafish, leptin was unable to improve glucose tolerance with levels identical to the control group (figure 3E, H). LiCl treatment improved glucose tolerance in all groups (22% for *lepa*^*nz301*^ fish, 15% for *lepb*^*nz302*^ fish, and 28% for *lepr*^*nz303*^ fish), demonstrating that a functional leptin receptor is not required for the glucose lowering effect of LiCl. No additive effect was found between leptin and LiCl.

**Figure 3.**
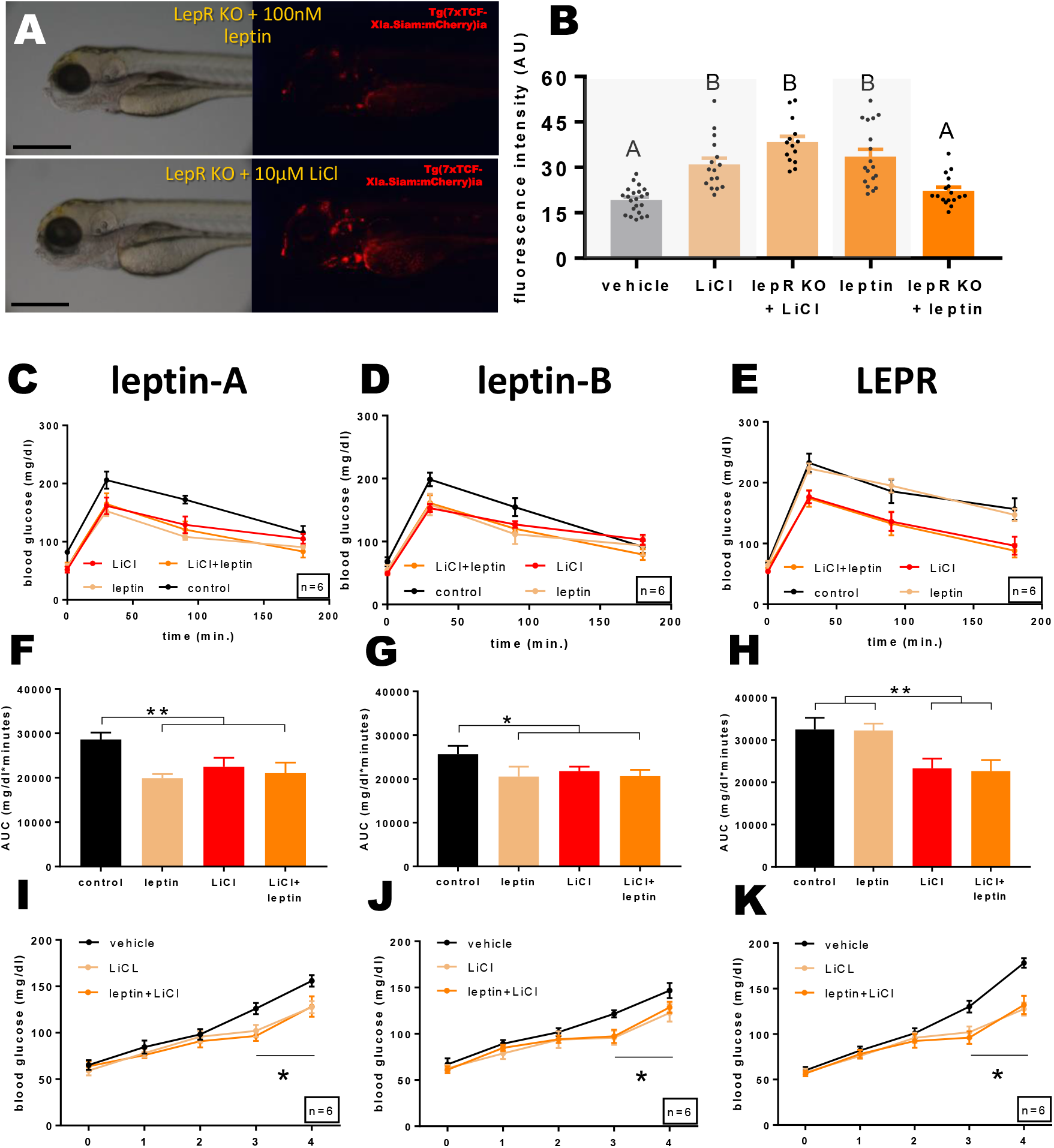
Activation of canonical Wnt signalling improves glucose tolerance in lepa^nz301^fish,lepb^nz302^ fish and lepr^nz303^zebrafish. (A) 5 dpf CRISPR-mediated leptin receptor-deficient Tg(7xTCFXla.Siam:nlsm Cherry)^ia5^ larvae treated with 100 nM recombinant leptin or 10 μM LiCl for 2 hours. Scale bar = 500 μM. (B) Fluorescence intensity in the hypothalamus of differentially treated 5 dpf Tg(7xTCF-Xla.Siam:nlsmCherry)^ia5^ larvae. A-B=P<0.05, one-way ANOVA. (C) Glucose tolerance of adult male lepa^nz301^ zebrafish (n=6). Fish were treated with 10 μM LiCl (three hours before glucose injection), with 0.6 g/L of recombinant mouse leptin dissolved in Cortland salt solution (one hour before glucose injection), with vehicle only, or with a combination of LiCl and leptin. Following 0.5 mg/g glucose injection, blood samples were taken at 30, 90, and 180 minutes post injection. (D) Same as (C), but for lepb^nz302^ fish. (E) Same as (C) and (B), but for lepr^nz303^ fish. (F) Area under the curve of (C). **P<0.001, one-way ANOVA. (G) Area under the curve of (D). *P<0.05, one-way ANOVA. (H) Area under the curve of (E). **P<0.001, one-way ANOVA. (I) Blood glucose values of adult male lepa^nz301^ zebrafish (n=6) over the course of a 4-day immersion in a 1% glucose solution. On the third day, fish were exposed to 10 μM LiCl for three hours before daily blood sampling. One hour before blood sampling, fish were injected intraperitoneally with 0.6 g/L of recombinant mouse leptin dissolved in Cortland salt solution, or with vehicle only. *P<0.05, repeated measures ANOVA. (J) Same as (G), but for lepb^nz302^ fish. *P<0.05, repeated measures ANOVA. (K) Same as (G) and (H), but for lepr^nz303^ fish. *P<0.05, repeated measures ANOVA.

In accordance, LiCl treatment attenuated persistent hyperglycemia in *lepa*^*nz301*^ fish (126.0±6.1 vs 102.0±6.5 mg/dl; figure 3 I), *lepb*^*nz302*^ fish (121.5±3.8 vs 95.7±7.8 mg/dl; figure 3 J) and *lepr*^*nz303*^ zebrafish (130.1±6.6 vs 102.0±6.5 mg/dl; figure 3 K). The effect of LiCl on blood glucose levels appears to be longer lasting than the effect of leptin (figure 2). LiCl-treated fish had significantly lower blood glucose levels not only immediately after the treatment ended, but on the following day as well (155.8±6.1 vs 127.5±6.7 mg/dl for *lepa*^*nz301*^ fish; 146.7±8.2 vs 122.5±9.4 mg/dl for *lepb*^*nz302*^ fish; 178.2±5.3 vs 127.5±7.2 mg/dl for *lepr*^*nz303*^ fish). Together, these data demonstrate that canonical Wnt pathway activation via LiCl-mediated inhibition of GSK3β regulates glucose homeostasis even in the absence of an intact leptin system.

### Inhibition of the canonical Wnt pathway blocks the glucoregulatory effect of leptin

To investigate whether the leptin-induced activation of Wnt is involved in mediating leptin action on glucose homeostasis, we pharmacologically activated or inhibited the canonical Wnt pathway, induced hyperglycemia acutely or persistently and then treated the fish with leptin. During an acute glycemic challenge in the form of an ipGTT, LiCl and leptin treatment reduced the AUC to a similar extent (∼20%), whereas combined application was not more effective (figure 4A, B). Under artificially-induced hyperglycemia (figure 4C, J), these effects were replicated with acute treatment with leptin, LiCl or both on day 3 of hyperglycemia resulting in a 10% reduction (LiCl), and a 15% reduction (leptin and leptin+LiCl) in glucose levels. To test whether the glucose-lowering effect of leptin was dependent on Wnt pathway activation, we applied PNU74654 two hours before acute leptin treatment. PNU74654 pretreatment led to a return of glucose levels to that observed in control conditions (figure 4D,E). PNU74654 alone, on the other hand, led to a 28% increase in AUC (figure 4,DE). During persistent hyperglycemia, Wnt pathway inhibition with PNU74654 did not significantly aggravate the rise of blood glucose (117±5.7 mg/dl) when compared with control fish (113±4.1 mg/dl). More importantly however, PNU74654 prevented the ability of leptin to lower blood glucose levels (107±5.3 mg/dl), compared with leptin-treated fish (97±3.1 mg/dl; *P<0.05, repeated measures ANOVA; figure 4F,J). Likewise, the anthelmintic drug pyrvinium pamoate impaired glucose tolerance by 39%, and blocked the glucose-lowering effect of acute leptin treatment in an ipGTT (figure 4G,H). Under artificially-induced hyperglycemia, pyrvinium did not significantly aggravate the rise of blood glucose (119±6.7 mg/dl) when compared with control fish (113±4.1 mg/dl), but prevented the ability of leptin to lower blood glucose levels (107±7.9 mg/dl), compared with leptin-treated fish (97±3.1 mg/dl; P<0.05; figure 4I,J). These findings demonstrate that intact canonical Wnt signalling is required for the ability of leptin to regulate blood glucose levels.

**Figure 4.**
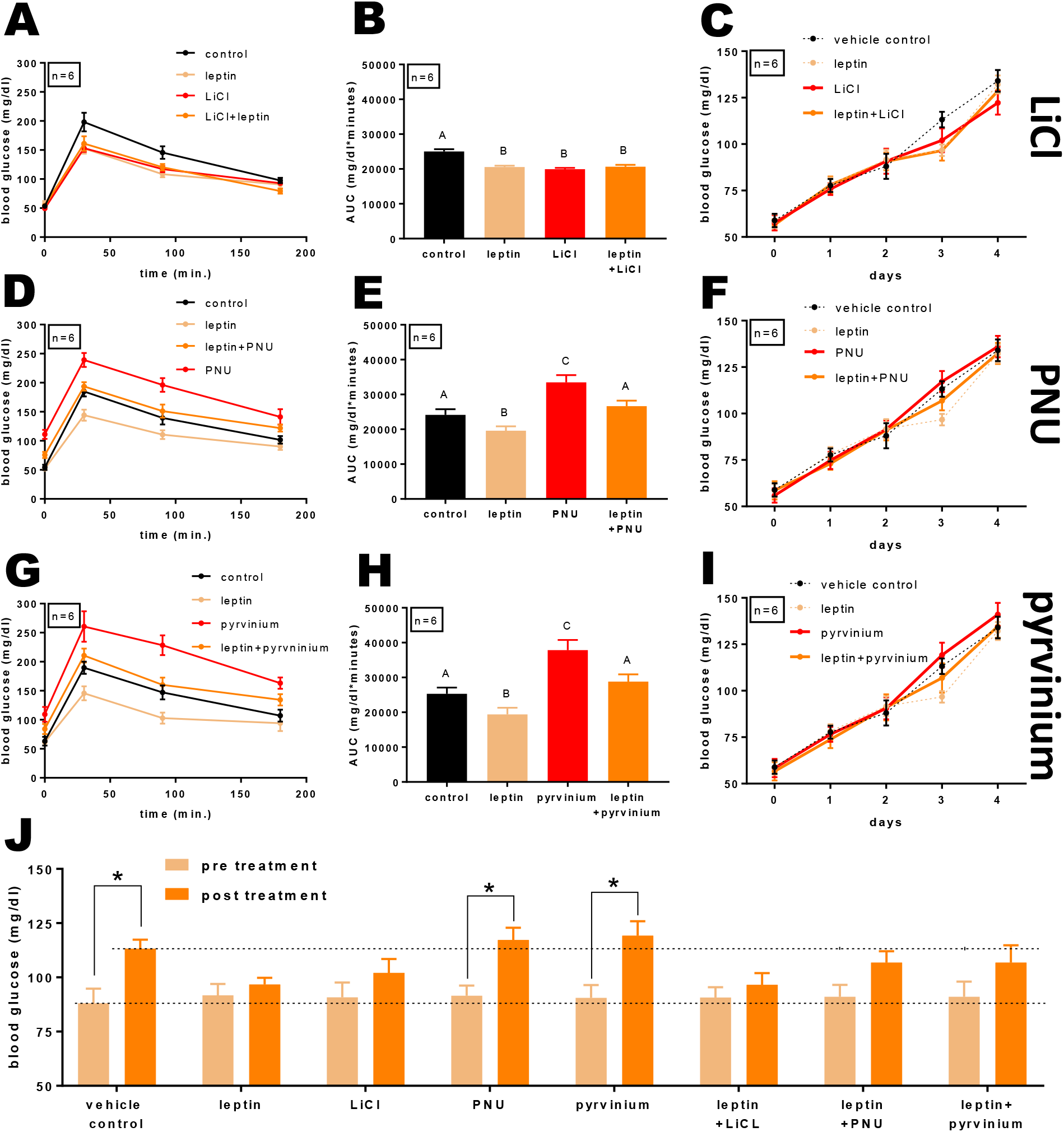
Glucose tolerance in wild type zebrafish following WNT pathway manipulation and leptin treatment. (A) Glucose tolerance of adult wild type male zebrafish (n=6). Fish were treated 10μM LiCl (three hours before glucose injection), 0.6 g/L of recombinant mouse leptin dissolved in Cortland salt solution (one hour before glucose injection), vehicle only, or a combination of LiCl and leptin. Following 0.5 mg/g glucose injection, blood samples were taken at 30, 90, and 180 minutes post injection. (B) Area under the curve of (A). A-B=P<0.05, one-way ANOVA. (C) Blood glucose values of adult wild type male zebrafish (n=6) over the course of a 4-day immersion in a 1% glucose solution. On the third day, fish were exposed to 10 μM LiCl for three hours before daily blood sampling. One hour before blood sampling, fish were injected intraperitoneally with 0.6 g/L of recombinant mouse leptin dissolved in Cortland salt solution, or with vehicle only. (D) Glucose tolerance of adult wild type male zebrafish (n=6). Fish were treated 10 μM PNU74654 (three hours before glucose injection), 0.6 g/L of recombinant mouse leptin dissolved in Cortland salt solution (one hour before glucose injection), vehicle only, or a combination of PNU74654 and leptin. (E) Area under the curve of (D). A-B and A-C=P<0.05, one-way ANOVA. (F) Blood glucose values of adult wild type male zebrafish (n=6) over the course of a 4-day immersion in a 1% glucose solution with exposure to 10 μM PNU74654. (G) Glucose tolerance of adult wild type male zebrafish (n=6). Fish were treated 10 μM pyrvinium pamoate (three hours before glucose injection), 0.6 g/L of recombinant mouse leptin dissolved in Cortland salt solution (one hour before glucose injection), vehicle only, or a combination of pyrvinium pamoate and leptin. (H) Area under the curve of (G). A-B and A-C=P<0.05, one-way ANOVA. (I) Blood glucose values of adult wild type male zebrafish (n=6) over the course of a 4-day immersion in a 1% glucose solution with exposure to 10 μM pyrvinium pamoate. (J) Comparison of blood glucose levels in (C), (F) and (I), pre-treatment (day 2) and post treatment (day 3). *P<0.05, repeated measures ANOVA.

### Conditional deletion of β-catenin in the mediobasal hypothalamus of male mice exacerbates DIO-induced body weight gain, food intake and causes leptin resistance

The data in zebrafish clearly showed that leptin activated Wnt signalling and that this was important for the effect of leptin on glucose homeostasis. Next, we sought to evaluate whether this mechanism could also be demonstrated in a mammalian model. We used a conditional deletion of β-catenin to ablate canonical Wnt signaling in the mediobasal hypothalamus (MBH). Global knockout of β-catenin is embryonic lethal [36], and hence, we employed β-catenin^*flox*^ mice and bilaterally injected AAV2-mCherry-iCre into the mediobasal hypothalamus to ablate β-catenin in the adult brain (β-catenin KO). We measured body weight and food intake daily following the introduction of AAV-iCre into the mediobasal hypothalamus until the conclusion of the experiment, and measured glucose homeostasis using a GTT. There was no significant difference in body weight or caloric intake between the control- and β-catenin KO mice when mice were fed low fat (control) diet (LFD) except between days 18-22 when we observed a mild increase in body weight (figure 5 A,B). However, mice lacking β-catenin in the MBH had markedly impaired glucose tolerance relative to controls (Figure 5 C, D; P<0.05). After 4 weeks fed LFD, both groups of mice were fed a high fat diet (HFD) for 6 weeks, which led to an increase in body weight irrespective of genotype. β-catenin KO mice, however, exhibited a much larger increase in body weight in response to HFD than mice injected with the control virus. This effect became significant from the 10^th^ day of HFD feeding (Figure 5 A, P<0.01). By the end of the study the β-catenin KO mice were 15% heavier than mice that received control injection and were fed HFD (P<0.0001).

**Figure 5.**
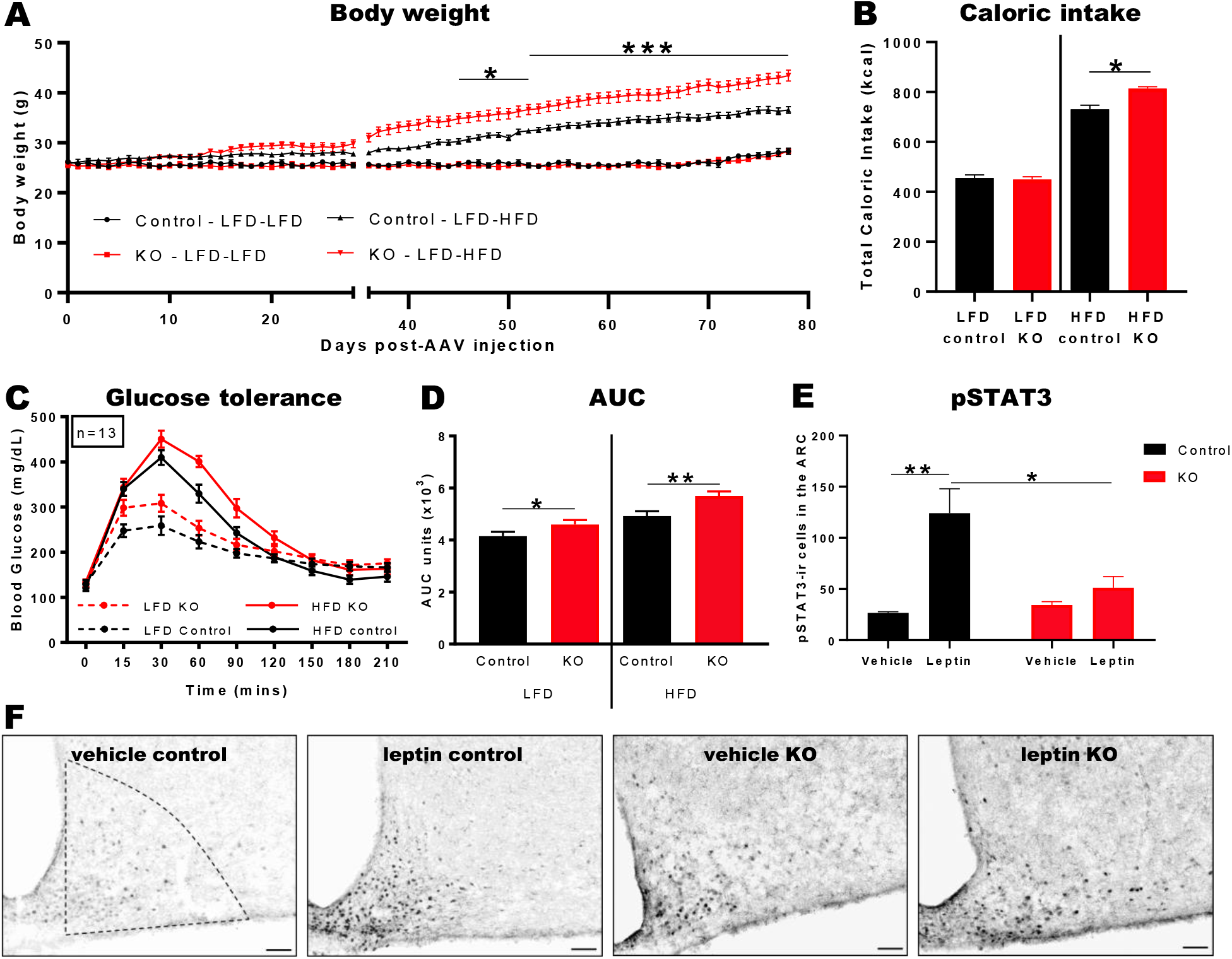
Conditional deletion of β-catenin in the mediobasal hypothalamus of male mice exacerbates DIO-induced body weight gain, food intake and causes leptin resistance. (A) Body weight of male β-catenin^flox^ mice injected with either the control or inducible Cre expressing virus maintained on LFD (ad libitum) for 4 weeks followed by 6 weeks on HFD (ad libitum). Also shown are the body weight (g) of β-catenin flox mice maintained on LFD throughout the experiment. The bar graphs represent total caloric intake (kcal) of male mice while on LFD and HFD. Error bars denote SEM. *, P<0.05; ***, P<0.001; KO compared to control mice. (B) Food intake of mice in (A). (C) Glucose tolerance of β-catenin KO- and control mice during intraperitoneal glucose tolerance tests. Glucose tolerance tests were performed 4 weeks after virus injection but maintained on LFD and 6 weeks after switching to HFD (B,E). (D) Area under the curve of (C). Error bars denote SEM. *, P<0.05; **, P<0.01; KO compared to control mice. (E) Quantified pSTAT3-immunoreactive positive cells in the arcuate nucleus after 30 mins of i.p. leptin (1.25 mg/kg) or PBS (vehicle) treatment for the different groups. Error bars denote SEM. *, P<0.05; **, P<0.01; leptin-compared to vehicle-treated mice at each group. (F) Representative micrographs of arcuate sections showing pSTAT3 expression following PBS or leptin treatment. Scale bar = 100 μM.

We also observed that during HFD feeding, the β-catenin KO animals had a higher caloric intake compared with the control mice (Figure 5 B, P<0.05). The impaired glucose tolerance induced by the lack of β-catenin in the MBH persisted after 6 weeks of HFD feeding (Figure 5 C, D; P<0.01) relative to controls.

We next assessed whether β-catenin deletion affects molecular leptin sensitivity by performing immunohistochemistry for pSTAT3 in the arcuate nucleus. As expected, acute injection of leptin (1.25 mg/kg body weight) induced a marked increase in pSTAT3 positive cells in control mice compared with vehicle-treated mice. In β-catenin mice, however, the response to leptin was fully ablated (P<0.05), suggesting the establishment of resistance to exogenously applied leptin (Figure 5 E, F; P<0.01 and P<0.05, respectively).

## Discussion

In this study we combined visualization of Wnt pathway activation in larval zebrafish brain with conditional genetic ablation of the Wnt pathway in adult mice to show: 1) Leptin induces activation of the Wnt pathway in the hypothalamus; 2) this action is dependent on the leptin receptor; 3) that activation of Wnt is sufficient to mimic leptin action on glucose homeostasis; 4) that leptin action on glucose homeostasis is impaired following pharmacological blockade of the Wnt pathway. Collectively, these data support the hypothesis that leptin regulates glucose homeostasis via the canonical Wnt pathway. These data provide new insights into why genetic polymorphisms in the Wnt pathway associate with increased risk of type 2 diabetes.

While raising zebrafish, we took great care to prevent any tank density effects on body weight, which have been shown to affect postembryonic development, somatic growth and fat accumulation [37]. Previous studies on teleost leptin knockout models have yielded contradicting results, with one study convincingly disproving a role for leptin as an adipostat in the zebrafish [30], whereas others reported an effect of leptin on adipostasis in zebrafish [38] and medaka [39]. Our data support the original observation [30], but extend this to also characterize a role for leptin on growth and body weight under conditions of caloric excess.

Glucose immersion of zebrafish has previously been shown to readily induce hyperglycemia over time [35]. We found that blood glucose levels in zebrafish were significantly elevated after two days of immersion, and that leptin administration on the third day consistently reduced glucose levels in wild type, *lepa*^*nz301*^ and *lepb*^*nz302*^, but not in *lepr*^*nz303*^ fish, confirming that the glucose-lowering properties of leptin are mediated by the leptin receptor. After 3 days of glucose immersion, blood glucose levels were higher in treated *lepr*^*nz303*^ fish than in *lepa*^*nz301*^ and *lepb*^*nz302*^as well as wild type fish. The elevated glucose levels in *lepr*^*nz303*^ compared with *lepa*^*nz301*^ and *lepb*^*nz302*^ fish suggests that the two leptin paralogs are probably functionally redundant for glucose regulation, and that only leptin receptor knockout is sufficient to induce hyperglycaemia. The fact that recombinant mouse leptin was active in zebrafish suggests that leptin function is highly conserved between species. This finding is in line with other studies that previously demonstrated anorexigenic effects of recombinant leptin in trout [40] and goldfish [41]. Interestingly, both the rate at which hyperglycemia was induced, and the potency of leptin to reduce hyperglycemia was identical in male and females. In humans, circulating leptin levels are higher in females [42], and the brains of female rats are more sensitive to the catabolic actions of ICV injected leptin than those of age- and weight-matched males [43]. Our data suggest that in the zebrafish, leptin acts sexually monomorphic.

It has been shown that zebrafish become obese when they are exposed to an overfeeding regime, and they display metabolic alterations similar to DIO mammals, like hypertriglyceridemia, hepatic steatosis, and systemic inflammation [44]. Under normal feeding conditions, knockout of the leptin receptor impaired glucose tolerance but had no effect on body weight regulation in the zebrafish, as neither knockout of leptin-a or leptin-b did. However, this was limited to fish fed normally. Overfeeding revealed an effect of leptin on body weight and standard length. Under these conditions *lepr*^*nz303*^ fish had elevated body weight compared with *lepb*^*nz302*^ fish and wild type zebrafish. Interestingly, *lepa*^*nz301*^ fish too also showed elevated body weight and standard length, suggesting a specific body weight regulatory and somatic effect of leptin-a that could not be compensated for by leptin-b. *In silico* binding simulation of zebrafish leptin-a and leptin-b predicts significantly lower binding energy to the leptin receptor for leptin-b [45]. Previous studies point towards a role for leptin-b in tissue regeneration rather than energy homeostasis [46, 47]. Further studies are required to delineate potential functional differences in downstream signal transduction between two leptin paralogues.

Overfeeding *per se* led to glucose intolerance in fish independent of genotype, and loss of leptin function in addition to overfeeding did not impair glucose tolerance further. One recent study found that overfeeding of zebrafish larvae leads to leptin resistance and reduced hypothalamic *pomca* levels, leading to activation of the melanocortin system, elevation of growth hormone levels, and enhanced somatic growth [48]. Together, these data point towards a fundamentally differential physiological role for leptin depending on nutrient availability. Under normal feeding conditions, leptin regulates glucose homeostasis. In times of nutrient excess on the other hand, leptin appears to regulate body weight and somatic growth. From an evolutionary perspective, this suggests that leptin originated as a glucoregulatory hormone, and that its adipostatic function in mammals may have been acquired at some point during evolution. Most aquatic species continue to grow somatically throughout life, whereas growth in terrestrial animals usually reaches a plateau due to gravity limitations. Because somatic growth limits movement much less in the water than on land, an adipostatic role of leptin may not be as crucial as in terrestrial species. This is in line with the recent discovery of the gravitostat in mammals [49]. This system has been suggested to regulate fat mass in obese mice independently of leptin, whereas leptin-mediated regulation of fat mass seems to be limited to healthy lean mice [50].

The canonical Wnt pathway has been shown to be activated by glucose in pancreatic β-cells, adipocytes, muscle cells and a macrophage cell line [51]. In mice, we demonstrated that Wnt signalling in the hypothalamus is impaired during obesity [24]. In the present study, we provide the first *in vivo* evidence of canonical Wnt pathway activation in the hypothalamus by leptin. Using LiCl as a positive control, we found that leptin-induced Wnt activation was especially prominent in the hypothalamic region in the brain of zebrafish larvae. Intriguingly, CRISPR-mediated knockout of the leptin receptor totally abolished this activation, suggesting that leptin activation of the Wnt pathway is solely mediated by the leptin receptor. Leptin receptor expression in the zebrafish is found not only in the hypothalamus, but also in a variety of peripheral organs, including the eye, gut, liver, pancreas, and heart [52]. Another region that showed high intensity of fluorescence after Wnt activation by LiCl was the heart. However, leptin did not significantly induce Wnt reporter-driven fluorescence in the heart.

Inhibition of the Wnt pathway blocks the ability of leptin to lower blood glucose levels both during acute and persistent hyperglycemia, suggesting that leptin regulates glucose homeostasis predominantly via the Wnt pathway. An antidiabetic action upon activation of the Wnt pathway has been confirmed for LiCl treatment, which has been shown to attenuate non-fasting blood glucose levels in diabetic Lep^ob/ob^ BTBR T+ Itpr3tf/J (BTBR) mice [53]. In accordance, LiCl treatment improves glucose tolerance and normalizes blood glucose levels during a persistent hyperglycemic challenge in zebrafish. Activation of the Wnt pathway with LiCl improved glucose homeostasis, even in leptin- or leptin receptor deficient fish, suggesting that LiCl acts independently of leptin and that leptin acts upstream of the canonical Wnt signalling cascade. We could previously show that leptin induces phosphorylation of LRP6 in the arcuate nucleus of the Djungarian hamster (*Phodopus sungorus*) [54].

The ability of Wnt signalling to regulate blood glucose levels is often ascribed to GSK3β being a site of convergence between canonical Wnt- and insulin signalling [55]. In a previous study we showed that neuron-specific overexpression of GSK3β in the hypothalamus exacerbated the effects of diet-induced obesity in wild type mice compared with mice fed a standard diet, measured as increased hyperphagia, obesity and glucose intolerance [25]. We also found that intracebroventricular injection of a GSK3β inhibitor or the WNT pathway antagonist Dickkopf 1 led to very rapid improvement or deterioration of glucose homeostasis, respectively. Here we inhibited the Wnt pathway both upstream (using pyrvinium pamoate) and downstream (using PNU74654) of GSK3β, yet both manipulations impaired glucose tolerance and blocked the glucoregulatory effect of leptin.

From these results it is unclear whether the glucoregulatory action of this pathway depends on transcriptional targets of the canonical Wnt pathway. Since β-catenin as a transcriptional coactivator of the pathway is crucial for activation of TCF7l2, we conditionally ablated β-catenin from the mediobasal hypothalamus in mice. Intriguingly, this treatment replicated the glucoregulatory effect observed in zebrafish, suggesting that the canonical Wnt pathway in the brain is a major player in the neuroendocrine regulation of whole body glucose homeostasis across different vertebrate species. This manipulation led to exogenous leptin resistance, confirming that leptin action largely depends on functional Wnt signalling in the hypothalamus.

Taken together we identify a novel essential role of the central canonical Wnt pathway in the neuroendocrine control of glucose homeostasis in zebrafish and mice. Furthermore, our findings highlight that leptin may primarily have evolved as a glucoregulatory hormone with its role of an adipostat acquired later in evolution. Finally, the glucoregulatory action of leptin is mediated via the Wnt pathway - an essential mechanism that appears to be conserved throughout the vertebrate phylum.

## Methods

### Ethics

Procedures involving animals were performed in accordance with national animal ethics legislation and received approval by University of Otago Animal Ethics Committee (AUP-18-121).

### Zebrafish Husbandry

Zebrafish (AB strain) were maintained in 3.5 L tanks on a Palletized Centralized Life Support System (Tecniplast). The water in this recirculating system was pumped through mechanical filtration, charcoal filtration, and UV-treatment; and 10% of the water was replaced every hour. The water was kept at 26–30°C, with pH 7.6–8.0 and a conductivity of 300–600 μS. The facility environment maintained a 14-hour light and 10-hour dark cycle. Water quality parameters were automatically measured and adjusted, and remained within acceptable limits for the duration of the study.

### CRISPR Cas9 mutagenesis

Single guide RNAs (sgRNAs) were synthesized *in vitro*. Cas9 mRNA was transcribed from a pT3TS-nCas9n plasmid (Addgene plasmid #46757). Offspring of AB or *Tg(7xTCFXla*.*Siam:nlsmCherry)*^*ia5*^ zebrafish were injected at the one cell stage into the cell with ∼1 nL of a solution containing zebrafish 212.2 ng/μL Cas9 mRNA and 35.4 ng/μL gRNA, based on [56]. As a positive control, and to test the quality of Cas9mRNA, we used an sgRNA targeting the tyrosinase gene. Mutagenic efficiency was analyzed using a three-primer fluorescence PCR method. Biallelic mutant founder fish (F0) were inbred, giving rise to stable mutant offspring. Target sequences were *lepr* GGAGCGCCAGTAAAGCCGTGTGG; *lepa* GGAATCTCTGGATAATGTCCTGG; *lepb* ACAGAACTGAGACCATCAATGGG; *tyr* GGACTGGAGGACTTCTGGGG.

### Zebrafish Overfeeding

3-month-old male leptin mutant fish and wild type control fish were assigned to either a 6-week overfeeding regime, consisting of 6 daily feeds, or a standard diet of 2 feeds per day. Feeds alternated between 20 mg/fish of ZM-400© fish pellets, and freshly hatched brine shrimp (*Artemia nauplii*, 30mg cysts/fish). ZM dry pellets (Zebrafish Management Ltd.) consisting of 58% protein, 14.5% fat, 11.5% ash, 7.0% moisture, 30,000 I.U./kg vitamin A, 2,500 I.U./kg vitamin D3, 400 mg/kg vitamin E, 2,000 mg/kg vitamin C, 30 mg/g ω3 highly unsaturated fatty acids. Feeding times were Zeitgeber Time (ZT) 1:00 (with lights turning on at ZT 0:00), ZT 3:30, ZT 5:00, ZT 7:30, ZT 9:00 and ZT 11:30h under the overfeeding regime, and ZT 3:30 and ZT 7:30 in the normal fed group. Feeding was done manually, and leftover food was removed by siphoning to prevent an effect of water quality on body weight [57]. Body weights were measured weekly. Standard length (SL), defined as the length measured from the tip of the snout to the posterior end of the last vertebra, was measured at week 0, week 3 and week 6. Finally, glucose tolerance was measured at the end of the dietary intervention

### Zebrafish Compound Exposure

Metformin (Sigma) was dissolved in fish water to a final concentration of 20 µM. The metformin solution was freshly prepared and changed daily. PNU74654 (Abcam) and pyrvinium pamoate (Sigma) were dissolved in DMSO and added to tank water or E3 medium at a final concentration of 10 µM. LiCl (Sigma) was dissolved directly in tank water or E3 medium at a concentration of 10 µM. *Tg(7xTCF-Xla*.*Siam:nlsmCherry)*^*ia5*^ were treated with 0.003% 1-phenyl-2-thiourea according to standard protocols to prevent pigmentation.

### Zebrafish Blood sampling

Borosilicate glass microcapillaries (Harvard Apparatus) were pulled on a Sutter p-97 Flaming Brown glass micropipette puller to create needles with a 1.0 mm outer diameter. Using scissors, the needle tips were cut obliquely to create a tip diameter of 100-300μm. Next, needles were heparinized (5mg/ml heparin in saline) using an aspirator tube assembly. For blood collection, a heparinized needle was inserted in the nosepiece end of the aspirator tube assembly. Adult zebrafish were anesthetized with 0.13% tricaine (3-aminobenzoic acid ethyl ether methanesulfonate, MS222). Anesthetized fish were carefully transferred onto soft tissue paper soaked in tricaine solution. Another soaked tissue was used to cover the fish’s head. The needle was then carefully inserted at a 30 – 45° angle into the dorsal aorta (DA), along the body axis and ventral to the spine. Generally, blood would rise into the needle in a pulsatile manner. If blood did not rise, gentle suction was applied through the mouthpiece, and the needle was moved gently by hand to encourage blood flow. The minimal required sample volume (0.6 μl) was collected. The needle was immediately removed, and gentle finger pressure with a soaked tissue was applied to the puncture site for ∼15 seconds or until bleeding stopped. Fish were then transferred to a recovery tank (28.5 °C), and water was gently swirled towards the gills.

### Zebrafish Glucose Immersion

The glucose immersion method was adapted from Capiotti et al. (2014). Fish were placed in standard housing tanks containing a 1% glucose solution (55.5 mM). Because the tanks were not on the normal recirculation system, solutions were renewed daily after feeding to prevent growth of microorganisms. Blood samples were taken daily.

### Zebrafish Intraperitoneal glucose tolerance tests

Fish were fasted for 72 hours to bring glucose levels down to baseline. Following anesthesia, fish were weighed and injected intraperitoneally with 0.5 mg glucose/g fish weight and allowed to recover for 30, 90, and 180 min after injection. Glucose concentrations were measured using a commercially available glucometer (Accu-Chek Performa; Roche)

### Mice

Mice containing lox-P sites in introns 1&6 of β-catenin (β-cateninflox; B6.129-Ctnnb1tm2Kem/KnwJ, Jackson labs; b-catenin gene is flanked by LoxP sites (floxed)) (8 weeks old; n=16-20 per group) were obtained from the University of Otago animal breeding facility. They were individually housed under 12:12h light/dark cycle (lights on 0600h) at a constant temperature (21 + 1 °C) with ad libitum access to food and water, except during fasting when only water was available. Mice were fed either a low-fat diet (LFD; D12450B Research Diets, New Brunswick, NJ 08901 USA) with 10% fat by energy (kcal) or high-fat diet (HFD; D12492 Research Diets) with 60% fat by energy (kcal). When required, in female mice the estrous cycle was determined by cytological examination of vaginal smears. This was done to ensure females were in diestrus when collecting tissue samples.

### Stereotaxic injections

Both adult male and female β-cateninflox mice were used. Intrahypothalamic injections were performed under isofluorane anaesthesia as described previously (23). Stereotaxic co-ordinates to reach the arcuate nucleus were 0.125mm posterior, ±0.35mm lateral and 0.59mm ventral relative to Bregma. An AAV-vector expressing Cre recombinase (AAV2-mCherry-iCre AAV virus, Penn Vector Core) was injected bilaterally using 1 mL Hamilton Syringes (Model 7001 KH SYR, 80100, Hamilton Company, Nevada 89502, USA) at volume of 0.5 mL at either side of the arcuate nucleus (Figure 1A). The injection needle was lowered to the correct coordinates over a period of 5 min, paused for 2 min, and then the virus was injected over a period of 2 min. The injection needle remained in place at each injection site for an additional 10 min to allow for diffusion and prevent backflow. The incision was then sutured and the mice were placed under a heating lamp in their home cage for recovery. The expectation was that viral-induced expression of the Cre would excise the floxed β-catenin gene resulting in a localized deletion of β-catenin at the site of injection. For control experiments, β-cateninflox mice were injected with an AAV-vector that did not express the Cre recombinase (AAV/DJ-CMV-mCherry AAV virus). For both knockout (KO) and control experiments, mice were body weight matched and placed in respective groups (control male mice, KO male mice; control female mice and KO female mice).

### Onset of obesity, food intake, metabolic measurement and glucose tolerance tests

Following the injection of the AAV-vector, mice were fed LFD for four weeks. Mice were fasted for 16 h, and at zeitgeber time (ZT) 0 (to guarantee a consistent influence of the circadian rhythm), they were injected intraperitoneally (ip) with glucose (1.5 g/kg), and a glucose tolerance test was performed. To determine the blood glucose levels, drops of blood from tail tips were collected repeatedly and glucose concentrations (mg/dL) were measured using a commercially available glucometer (Accu-Check Performa, Roche, Basel, Switzerland). For statistical validation, the area under the curve (AUC) was calculated.

We next investigated the effect of HFD on these mice. They were then fasted for 16 h and an ipGTT was performed as described above. Throughout the whole procedure, from viral injection till the conclusion of the study, daily body weight and food intake measurements were recorded. In addition, another cohort of mice was placed through a similar regime as above, but was fed LFD throughout the experiment.

### Immunohistochemistry – Validation of AAV-vector injections

To validate successful deletion of β-catenin in the arcuate nucleus, in another set of brain sections, immunohistochemical analysis of β-catenin (6B3; Rabbit mAb #9582; 1:100; Cell Signalling) was performed as per the manufacturers protocol. Images of the arcuate nucleus were taken, and presence or absence of β-catenin was analysed within that region.

### Immunohistochemistry – Onset of leptin resistance

To determine the onset of either leptin or insulin resistance in mice with β-catenin knocked out in β-catenin expressing cells specifically in the ARC, mice were body weight-matched and further subdivided into either vehicle- or leptin-or insulin-treated mice. At the conclusion of the study, mice were fasted for 16 h to reduce endogenous levels of either leptin or insulin, and at ZT0, were injected ip with either 0.1M PBS (vehicle), leptin (1.25 mg/kg; R&D Systems, Minneapolis, MN 55413 USA) or insulin (1 mg/kg; I6634; Sigma-Aldrich). Thirty mins (leptin-treatment) or 15 mins (insulin-treatment) post-injection, mice were anaesthetised (with Pentobarbitol; 30 mg/kg), and once the pedal withdrawal reflex was lost, were transcardially perfused with 0.9% heparinized saline followed by 4% paraformaldehyde in 0.1M phosphate buffer (pH 7.4). The procedure has been described in detail elsewhere (25).

The immunohistochemical analysis of phosphorylated signal transducer and activator of transcription3 (pSTAT3; Tyr705; Rabbit mAb #9145; 1:3000; Cell Signalling, Danvers, MA 01923, USA), a marker for leptin receptor signalling (25) or of phosphorylated Akt (Protein Kinase B) (pAkt; Ser 473; Rabbit mAb #4060; 1:500; Cell Signalling), a marker for insulin receptor signalling (26) was performed on 30 μm coronal brain sections in accordance with a previously described protocol for leptin (27) or insulin (26). pSTAT3 or pAkt positive immunoreactive cells were examined using an Olympus AX70 Provis light microscope (Olympus, Tokyo, Japan). Images of the arcuate nucleus were taken using the Spot RT Colour Camera attached to the microscope with an identical setting throughout the analysis. Two investigators who were blinded to the treatment counted immunoreactive cells within one of the bilateral halves of the arcuate nucleus (n=4-5 sections per mouse).

### Statistics

Data were analysed by one- or two-way ANOVA with repeated measurements tests, where appropriate followed by a Holm-Sidak post-hoc test to check for significance, as appropriate using Prism software. Results are presented as the mean + SEM. P<0.05 was considered statistically significant. For metabolic measurements, analysis of covariance (ANCOVA) for two independent samples was performed, whereby the body weight was used as the concomitant variable whose effects were brought under statistical control, and the energy expenditure was the dependent variable of interest. P<0.05 was considered statistically significant.

## Acknowledgements

This work was supported by the Royal Society of New Zealand Marsden Fund to AT, the Maurice Wilkins Centre (to A.T. and D.G) and the Health Research Council of New Zealand (to D.G.).

## Supplemental figures

**Figure S1.**
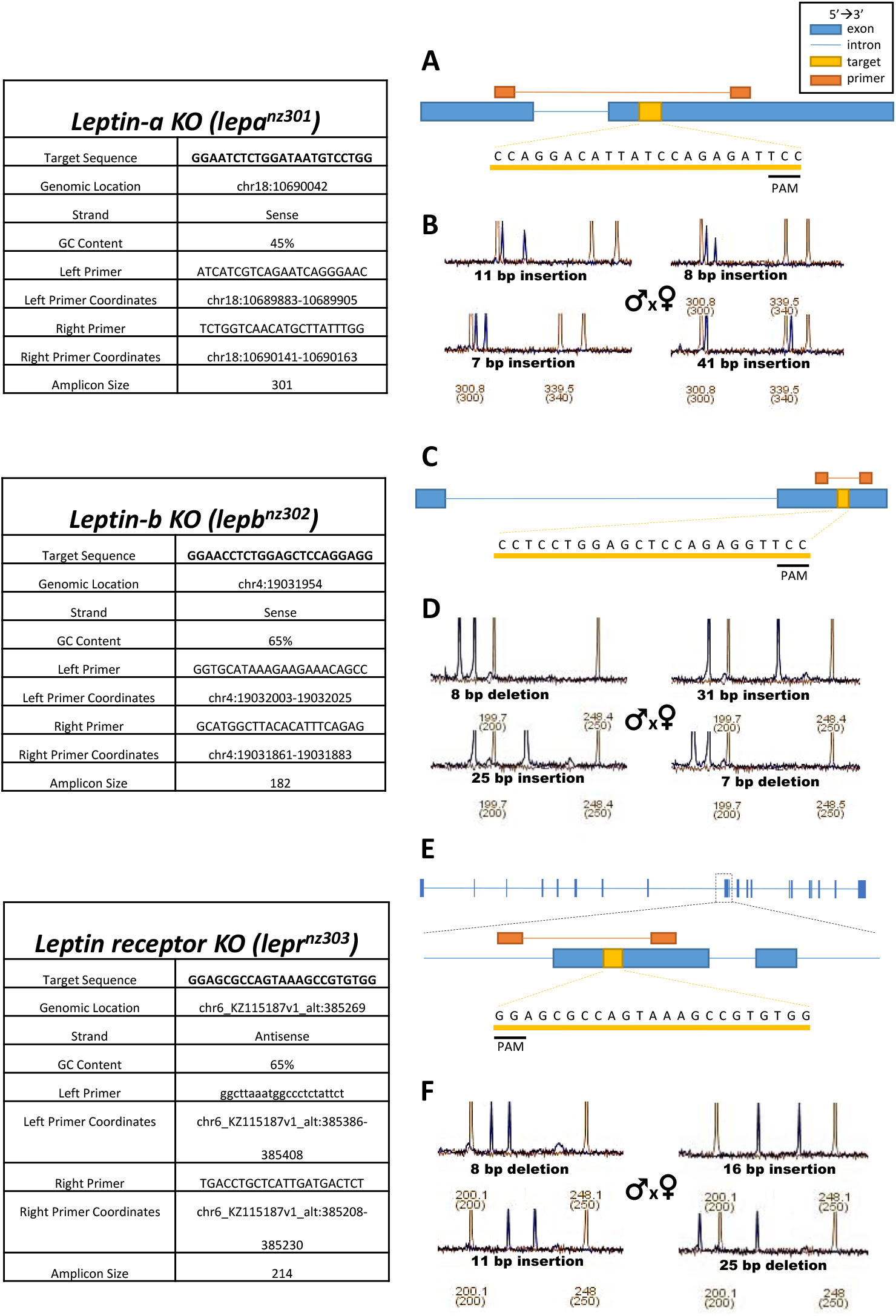
Creation of lepa^301^,lepb^302^,lepr^303^mutant zebrafish lines. (A) Zebrafish leptin-a gene with target sequence and primers. (B) Fluorescence PCR plots of selected lepa^301^ founder fish with annotated insertion/deletion. (C) Zebrafish leptin-b gene with target sequence and primers. (D) Fluorescence PCR plots of selected lepb^302^ founder fish with annotated insertion/deletion. (E) Zebrafish leptin receptor gene with target sequence and primers. (F) Fluorescence PCR plots of selected lepr^303^ founder fish with annotated insertion/deletion.

**Figure S2.**
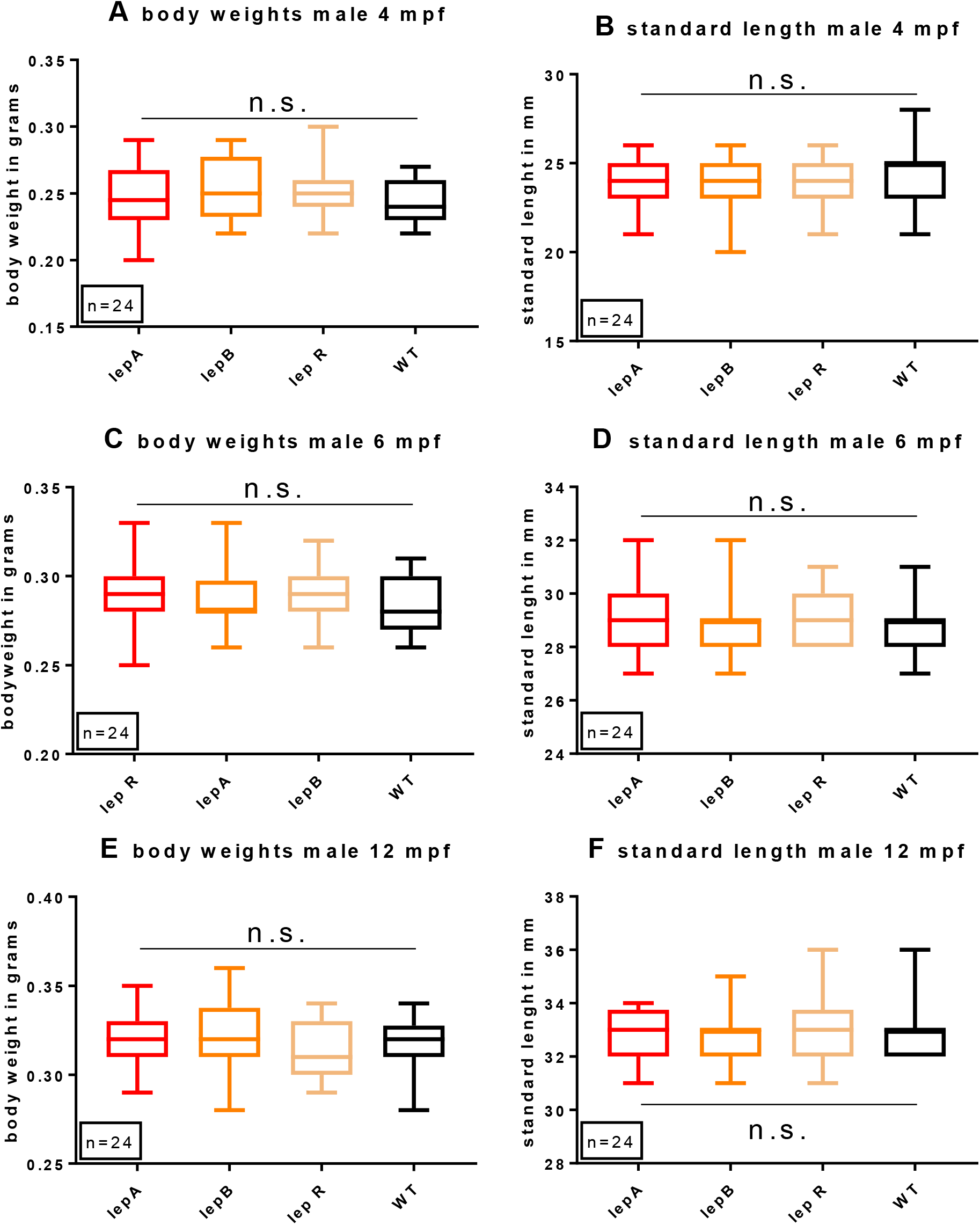
Leptin has no effect on male body weight or body length in the zebrafish. lepa^nz301^ fish, lepb^nz302^ fish, lepr^nz303^ fish and wild type controls (n=24) were raised at identical tank densities. Body weight was measured at 4 months’ post fertilization (A), 6 months post fertilization (C) and 12 months post fertilization (E). Standard length was measured simultaneously (B, D, F). No significant differences were found between any groups at any stage, as measured using one-way ANOVAs. lepA= lepa^nz301^ fish; lepB= lepb^nz302^ fish; lepR= lepr^nz303^ fish; WT=wild type control; data displayed as boxplots with whiskers representing minimum and maximum values.

**Figure S3.**
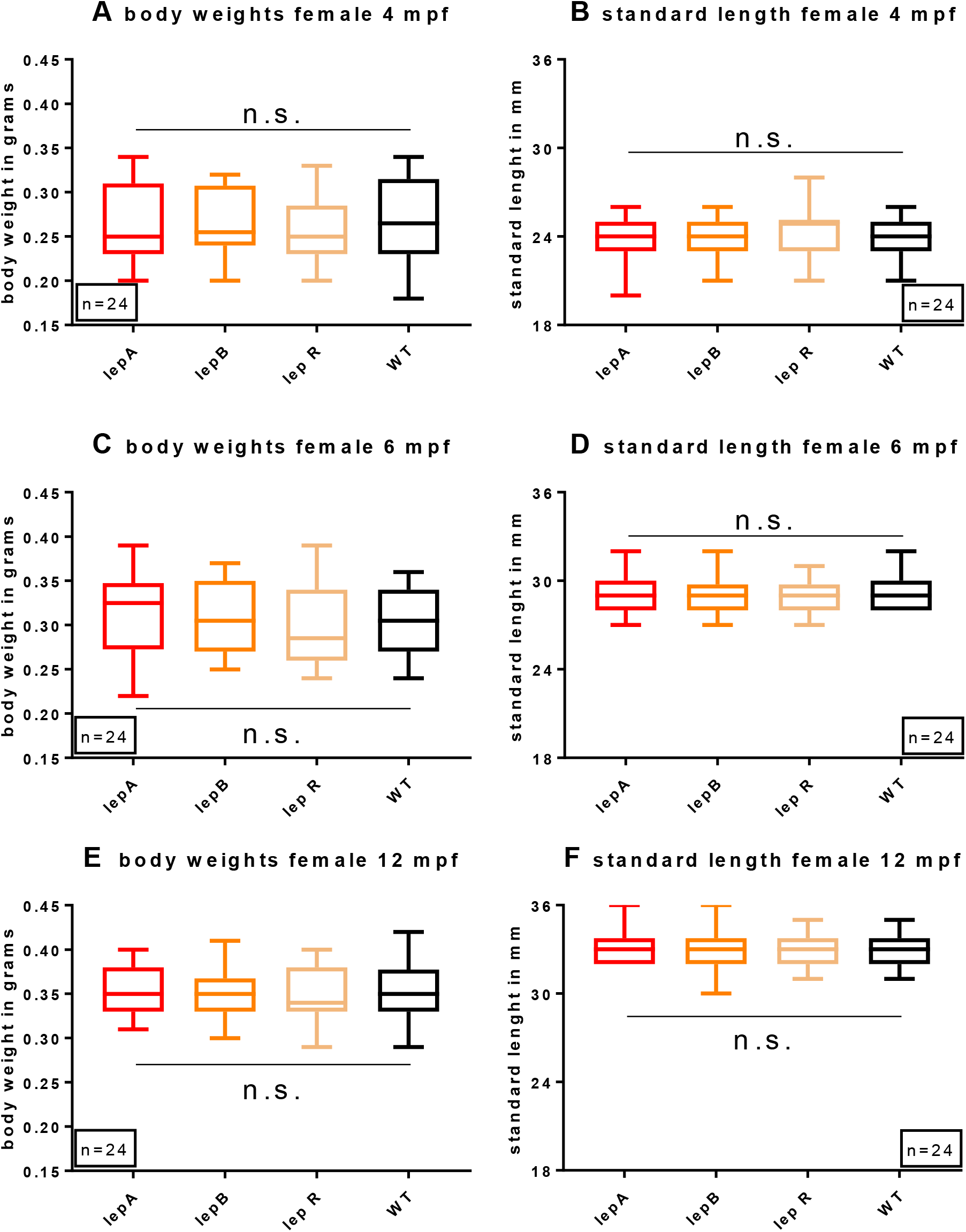
Leptin has no effect on female body weight or body length in the zebrafish. lepa^nz301^ fish, lepb^nz302^ fish, lepr^nz303^ fish and wild type controls (n=24) were raised at identical tank densities. Body weight was measured at 4 months post fertilization (A), 6 months post fertilization (C) and 12 months post fertilization (E). Standard length was measured simultaneously (B, D, F). No significant differences were found between any groups at any stage, as measured using one-way ANOVAs. lepA= lepa^nz301^ fish; lepB= lepb^nz302^ fish; lepR= lepr^nz303^ fish; WT=wild type control; data displayed as boxplots with whiskers representing minimum and maximum values.

**Figure S4.**
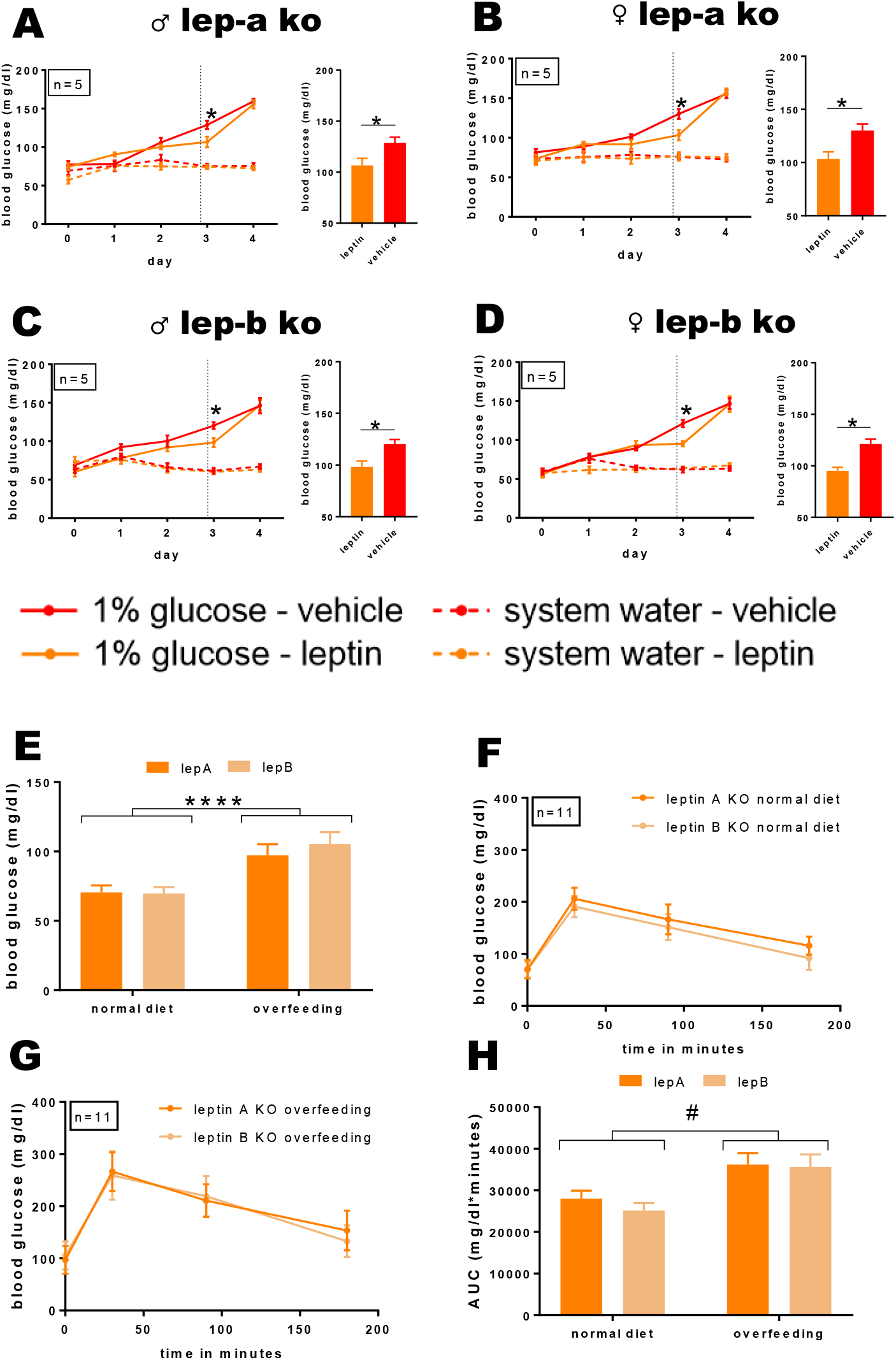
Regulation of glucose homeostasis and body weight in lepa^nz301^fish and lepb^nz302^fish. (A-D) Blood glucose values of male and female lepa^nz301^ fish and lepb^nz302^ fish over time following immersion in a 1% glucose solution. On the third day, one hour before blood sampling, fish were injected intraperitoneally with recombinant mouse leptin (2mg/kg) or vehicle. *P<0.05, repeated measures one-way ANOVA. Data displayed as mean±SEM (E) Baseline blood glucose levels of lepa^nz301^ fish and lepb^nz302^ fish. ****P<0.0001, two-way ANOVA. (F) Glucose tolerance in lepa^nz301^ fish and lepb^nz302^ fish (n=12). (G) Glucose tolerance in overfed lepa^nz301^ fish and lepb^nz302^ fish (n=12). (H) Area under the curve of (F) and (G). *P<0.05, one-way ANOVA; #P<0.05, two-way ANOVA. Data displayed as mean ± SEM.

**Figure S5.**
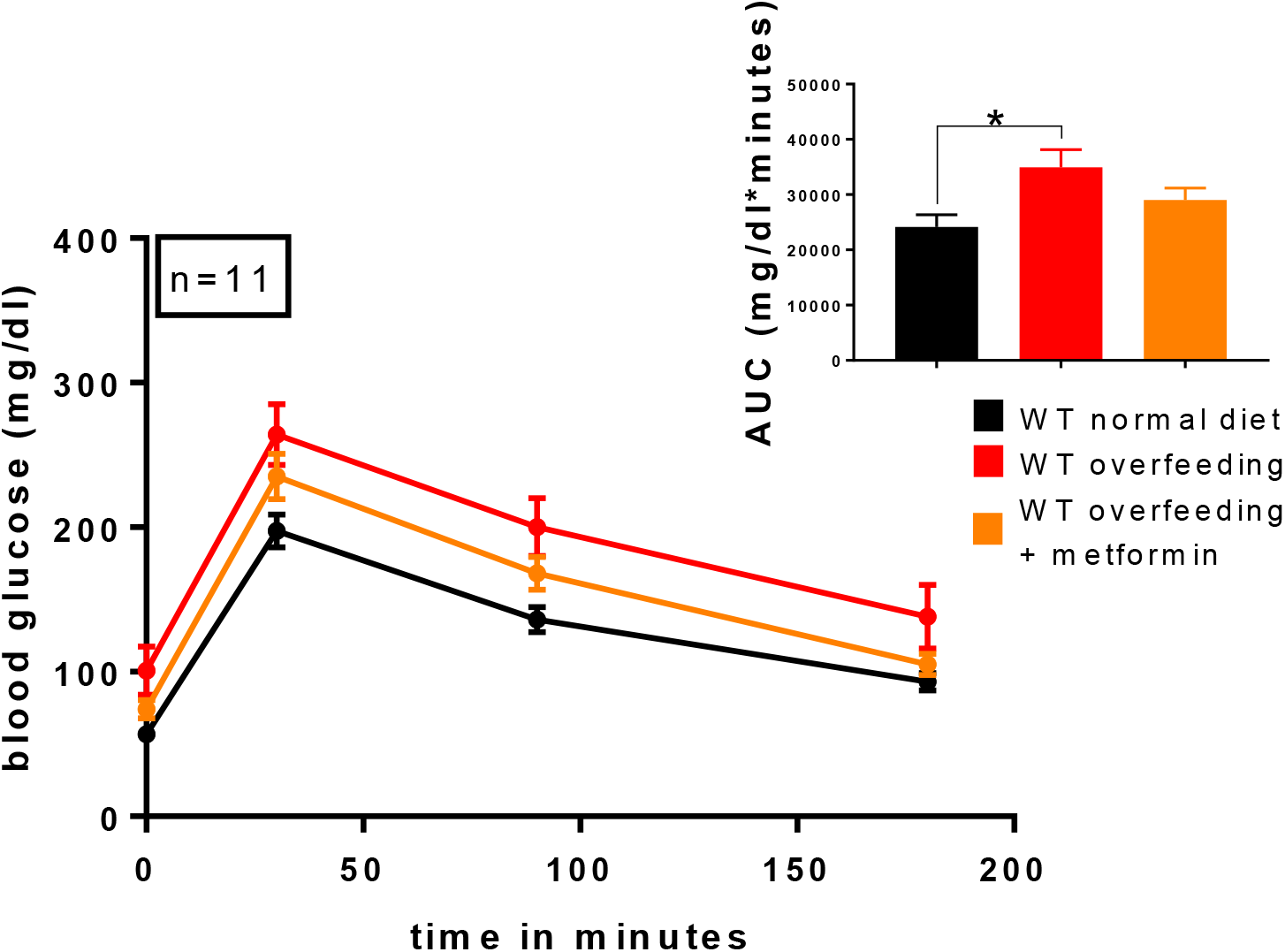
Metformin improves glucose tolerance in the zebrafish. After 5 weeks of overfeeding or normal diet, and one week of metformin (20 µM) treatment, wild type zebrafish (n=11) were challenged with a 0.5mg/g glucose solution, administered via IP injection. Blood samples were taken at 0, 30, 90 and 180 minutes. Overfed fish had significantly impaired glucose tolerance (34950±3211 mg/dl*minutes) compared to normally fed fish (24147±2192 mg/dl*minutes; *P<0.05, one-way ANOVA with multiple comparisons), but not compared to metformin-treated overfed fish (29033±2169 mg/dl*minutes). Main: blood glucose values over time; Insert: area under the curve of glucose tolerance test depicted in main figure. Data displayed as mean ± SEM.

**Figure S6.**
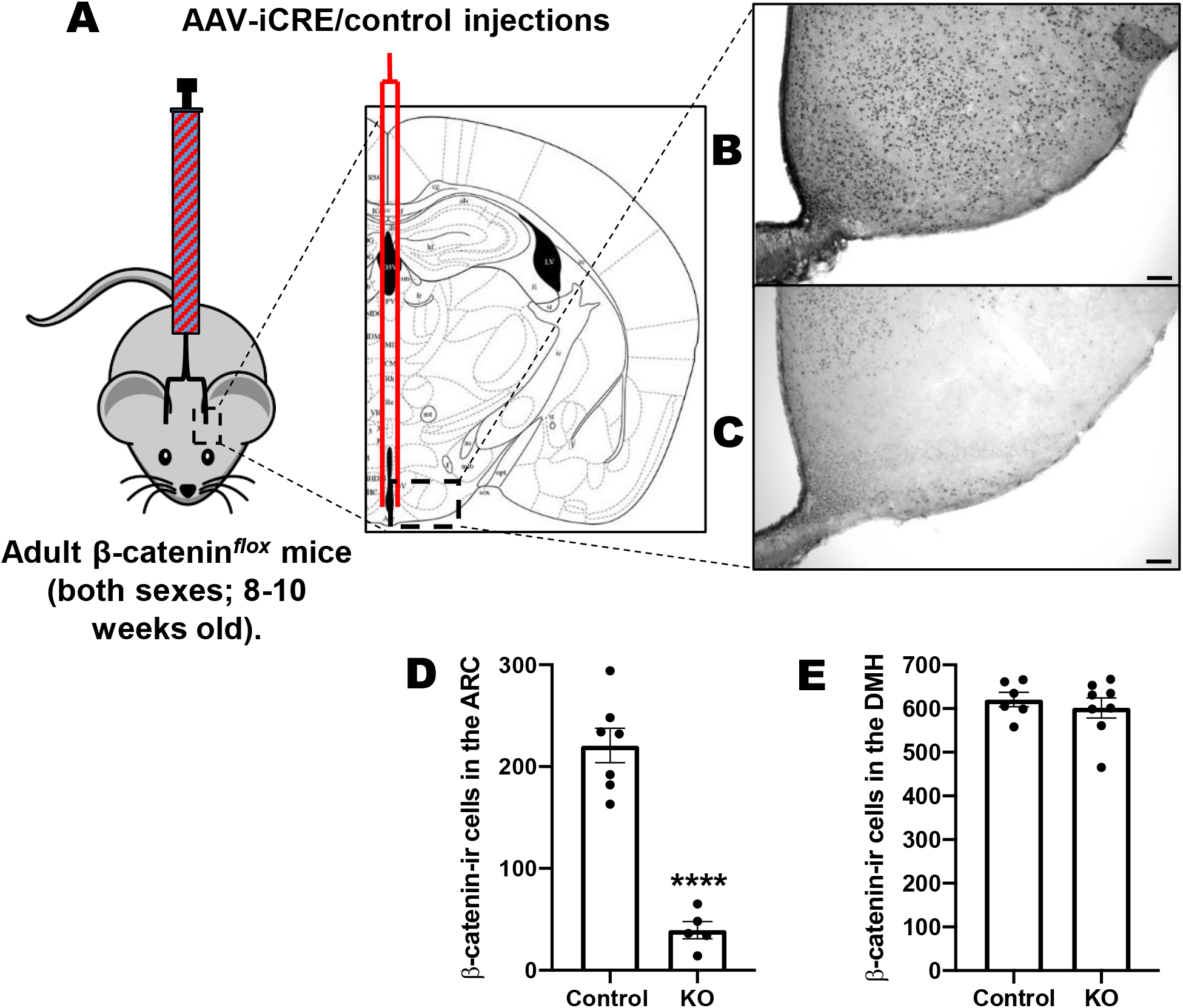
Validation of conditional β-catenin knockout in adult mice. (A) Overview of injection site. (B) Representative image of β-catenin staining in the mediobasal hypothalamus of a control-injected mouse. Scale bar = 100 μm. (C) Representative image of β-catenin staining in the mediobasal hypothalamus of an AAV-iCre-injected mouse. Scale bar = 100 μm (D) Quantification of β-catenin-positive cells in the arcuate nucleus of AAV-iCre-vs control - injected mice (****P<0.0001; Student’s t-test).(E) Quantification of β-catenin-positive cells in the dorsomedial hypothalamus of AAV-iCre-vs control-injected mice. No significant difference. Data displayed as mean ± SEM.

